# Plant Extracellular Vesicles Contain Diverse Small RNA Species and Are Enriched in 10 to 17 Nucleotide “Tiny” RNAs

**DOI:** 10.1101/472928

**Authors:** Patricia Baldrich, Brian D. Rutter, Hana Zandkarimi, Ram Podicheti, Blake C. Meyers, Roger W. Innes

## Abstract

Small RNAs (sRNAs) that are 21 to 24 nucleotides (nt) in length are found in most eukaryotic organisms and regulate numerous biological functions, including transposon silencing, development, reproduction, and stress responses, typically via control of the stability and/or translation of target mRNAs. Major classes of sRNAs in plants include microRNAs (miRNAs) and small interfering RNAs (siRNAs); sRNAs are known to travel as a silencing signal from cell to cell, root to shoot, and even between host and pathogen. In mammals, sRNAs are transported inside extracellular vesicles (EVs), which are mobile lipid compartments that participate in intercellular communication. In addition to sRNAs, EVs carry proteins, lipids, metabolites, and potentially other types of nucleic acids. Here we report that plant EVs also contain diverse species of sRNA. We found that specific miRNAs and siRNAs are preferentially loaded into plant EVs. We also report a previously overlooked class of “tiny RNAs” (10 to 17 nt) that are highly enriched in EVs. This new RNA category of unknown function has a broad and very diverse genome origin and might correspond to degradation products.

## INTRODUCTION

Small RNAs (sRNAs) are important 21 to 24 nucleotide (nt) non-coding signaling molecules involved in a wide variety of processes, including plant development, reproduction and defense (Samad et al., 2017). sRNAs can be divided into two categories, microRNAs (miRNAs) and small interfering RNAs (siRNAs), based on the differences in their biogenesis and mode of action. miRNAs originate from a single-stranded, self-complementary, non-coding RNA that forms a hairpin structure. In contrast, siRNAs originate from a double-stranded RNA molecule synthesized by RNA-DEPENDENT RNA POLYMERASES (RDRs). siRNAs can further be divided into two main categories, heterochromatic siRNAs (hc-siRNAs) and phased siRNAs, including *trans*-acting siRNAs (tasi-RNAs). Both of these double-stranded RNA structures are recognized by Dicer-like proteins (DCL), which cleave these RNAs into defined length products. One strand of these products is then selectively loaded onto an ARGONAUTE (AGO) protein and incorporated into the RNA-induced Silencing Complex (RISC). The RISC uses the sRNA in a sequence-homology-dependent manner to negatively regulate targets, typically mRNAs (Borges and Martienssen, 2015).

sRNAs are often mobile and function in non-cell autonomous silencing, which can be either local or systemic. Local RNA silencing occurs among groups of adjacent cells and can gradually spread from cell to cell (Marín-González and Suárez-López, 2012; Dunoyer et al., 2013). Systemic silencing occurs over long distances and can spread throughout an entire plant. While there are several documented cases of mobile small RNAs in plants, the mechanisms by which these RNAs move has yet to be clearly established. Local RNA silencing is thought to involve the transport of RNAs through plasmodesmata (PD) with the aid of PD-interacting proteins. The same is thought to be true for systemic silencing, although this process also likely requires transport of RNA into and through the phloem. Systemic RNA silencing progresses in a vascular-like pattern reminiscent of the phloem and is graft transmissible, moving bidirectionally up and down the plant in a manner that is heavily dependent on sink and source relationships (Lee and Frank, 2018).

Plant sRNAs are also known to move from plant cells into pathogens. Through a phenomenon known as Host-Induced Gene Silencing (HIGS), small RNAs produced in a plant cell can regulate the expression of genes in an invading pathogen or parasite. In recent years, HIGS has become a powerful tool for engineering resistance in crop plants to nematodes, insects, fungi, oomycetes and parasitic weeds (Cai et al., 2018). Although multiple studies have developed artificial HIGS systems, the phenomenon can occur naturally as well, such as in the transfer of microRNAs miR166 and miR159 from cotton into the hemibiotrophic fungus, *Verticillium dahliae* (Zhang et al., 2016), and the transfer of siRNAs from *Arabidopsis* into *Botrytis cinerea* (Cai et al., 2018). How this translocation is mediated is not well understood.

In mammals, extracellular vesicles (EVs) are known to mediate long distance transport of both coding and non-coding RNAs. RNAs protected within the lumen of an EV are transported to distant cells, where they retain their function after delivery (Ratajczak et al., 2006; Valadi et al., 2007; Mittelbrunn et al., 2011). The RNA contents of mammalian EVs largely reflects that of their cell type of origin, but can show enrichment for certain mRNAs and miRNAs. The overrepresentation of specific sequences inside EVs suggests an active mechanism for loading RNAs into these compartments. Usually, miRNAs make up a sizable percentage of the RNAs within EVs. However, other classes of small non-coding RNAs are also present, including tRNAs, rRNAs and fragments originating from both coding and non-coding RNAs. These other sRNAs are often highly enriched in EVs compared to their cell of origin. The exact role of other non-coding RNAs and RNA fragments in EVs has yet to be determined. They may function in signaling or they may represent waste products that EVs remove from the cell (Bellingham et al., 2012; Nolte’t Hoen et al., 2012; Huang et al., 2013; Chevillet et al., 2014; van Balkom et al., 2015).

Although the RNA content of mammalian EVs has been extensively characterized, the RNA content of plant EVs has not been carefully assessed. To address this knowledge gap, we optimized methods for purifying EVs from plant apoplastic wash fluid (Rutter and Innes, 2017), performed sRNA-seq analysis on EV RNA, and compared the resulting data to sRNAs in the source leaf tissue and EV-depleted aplastic wash fluid. These analyses revealed that plant EVs are enriched in single-stranded small RNAs, especially a class of RNAs ranging in size from 10 to 17 nts. These RNAs seem to represent degradation products of small RNA production. Their abundance in plant EVs raises the question of whether they serve a function in either intercellular and/or inter kingdom communication.

## RESULTS AND DISCUSSION

To analyze the sRNA content of EVs, we constructed sRNA libraries from three biological replicates of *Arabidopsis thaliana* rosette leaves (“total RNA”, henceforth), EVs, and apoplastic wash fluids depleted of EVs (“apoplast” henceforth), as previously described (Rutter and Innes 2017). After an initial step of standard read trimming (adaptor removal and size selection from 18 to 34 nt), we observed that only 20% of the reads from EV samples were retained. This very low rate of retention was due to a large proportion of very short reads, from 10 to 17 nt long, present in the EV samples. We named these unusually short reads “tiny RNAs” or tyRNAs. Changing the trimming criteria to retain any read from 10 to 34 nt allowed us to achieve an average of ~60% read retention (Supplemental Table 1). The size distribution of the EV and apoplast samples was unexpected (Figure 1A). Total RNA samples had the typical size distribution of *Arabidopsis* small RNAs, a peak at 24 nt followed by secondary peaks at 21 and 22 nt. However, EV samples had a high proportion of tyRNA reads between 10 and 15 nt and no peak in the 20 nt range, while apoplast samples had a high proportion of longer reads, between 30 and 32 nt (Figure 1A, left panel). When considering only distinct reads, where tags with the exact same sequence were counted only once (Figure 1A, right panel), EV samples continued to show an enrichment for tyRNAs. This suggests that the tyRNAs present in EVs were not only abundant, but also highly diverse in their sequence composition. Comparatively, apoplastic sRNAs were not enriched for tyRNA sequences. This strongly argues that tyRNA reads were not a result of degradation occurring during RNA isolation or library preparation, as these steps were performed in parallel for all RNA samples.

**Figure 1.**
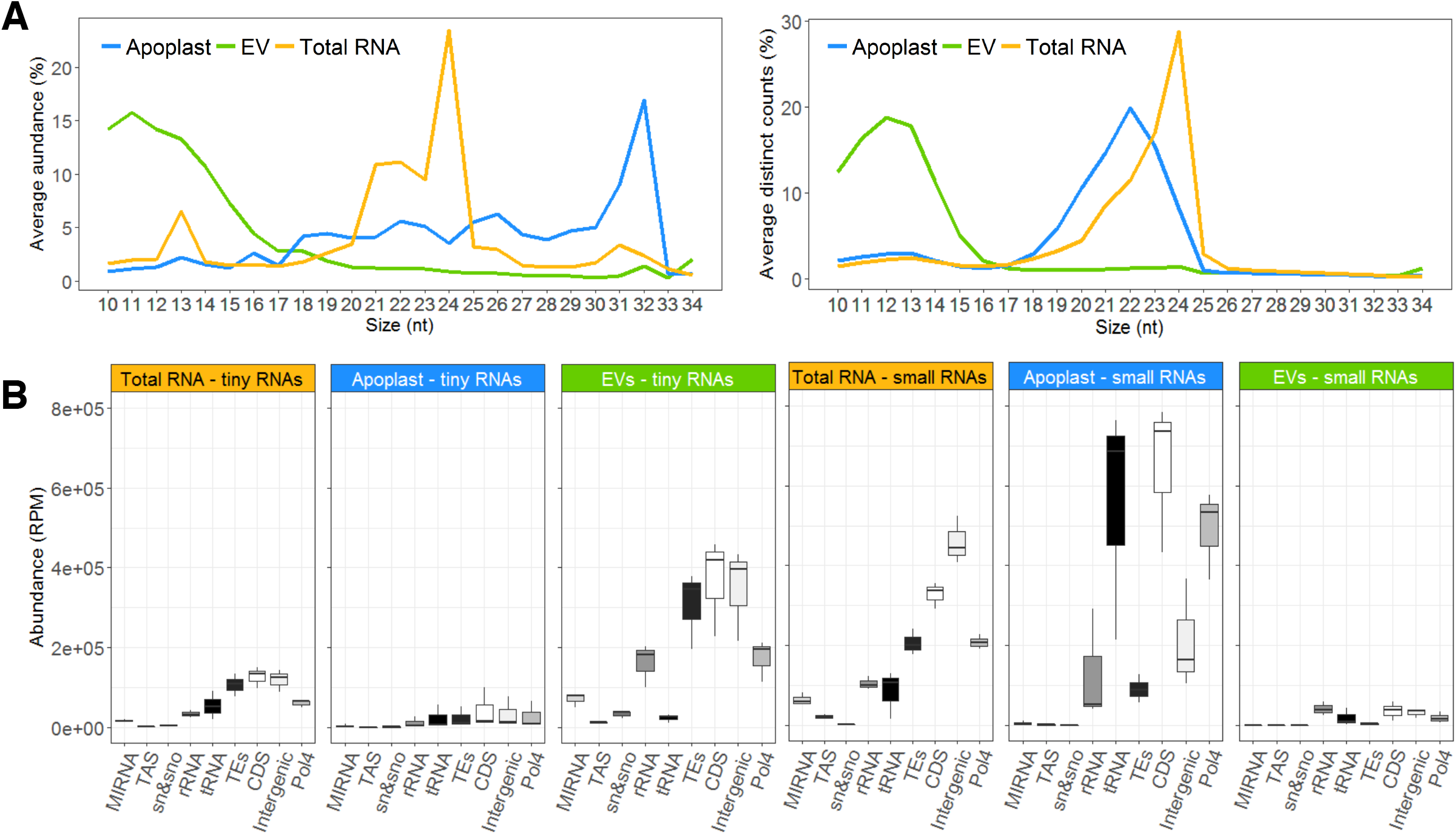
EVs are Enriched in Tiny RNA Sequences Derived from Diverse Sources. **(A)** Size distribution of sRNAs mapping to the genome; the percentage of each size class was calculated for each source of data, represented by the average of three replicate libraries. The x axis indicates the sRNA size and the y axis indicates its proportion. The panel at the left indicates the sRNA sizes and proportions, as measured by total abundance of the reads, and at right, as calculated by distinct reads (unique sequences). **(B)** The abundance of reads mapping to nine different features of the *Arabidopsis* genome. These include the following from left to right: *miRNA* precursors, transacting siRNAs (TAS) precursors, small nuclear RNAs (snRNA and snoRNAs); ribosomal RNAs; tRNAs; transposable elements (TEs); coding sequences (including introns and UTRs; CDS); intergenic sequences; and siRNA precursors dependent on RNA Polymerase IV (Pol4).

### Tiny RNAs mostly originate from CDSs, TEs and intergenic regions

To understand the genomic origin of these tyRNAs, we mapped these reads to different features of the *Arabidopsis* genome, including MIRNAs (genes encoding miRNA precursors), *TAS* genes (genes giving rise to tasiRNAs), transposable elements (“TEs”), rRNAs, tRNAs, small nuclear and small nucleolar RNAs (snRNAs and snoRNAs), coding sequences (“CDSs”), potential RNA Polymerase IV products (“Pol4”) (as defined in Blevins et al., 2015) and intergenic regions (Figure 1B). For reads that mapped to more than one feature, we counted each feature once. We performed the same analysis with the sRNA reads (18 to 34 nt) and observed that relative to sRNAs, tyRNAs were enriched in sequences that map to MIRNAs and TEs (Figure 1B). While the EV samples contained a higher abundance of tyRNA reads, tyRNAs from both total RNA and EV samples mapped to a similar distribution of features. In both samples, the majority of tyRNAs mapped back to TEs, CDSs and intergenic regions, which combined, comprise the largest proportion of the *Arabidopsis* genome (Figure 1B). We also detected high levels of tyRNAs derived from Pol4 precursors and rRNA regions, which represent 14% and 0.12% of the genome, respectively. Typically, rRNA represents around 80% of total leaf RNA, so the relatively low frequency of rRNA reads in each library indicates that they were not significantly contaminated with degraded cellular RNA. Together, these observations strongly suggest that EVs are highly enriched for tyRNAs and that this new RNA category might correspond to specific degradation products of sRNA precursors.

### MIRNA-derived tyRNAs are not randomly distributed

To determine whether tyRNAs are products of sRNA processing, we next analyzed their relative position in non-coding RNAs. The abundance of tyRNAs derived from miRNA precursors was almost twice that of snRNAs, snoRNAs, and tRNAs. Since this posits a clear preferential accumulation of MIRNA-derived tyRNAs in EVs, we analyzed the location of tyRNA sequences within miRNA precursors. In addition to a 5’ cap and a 3’ poly-A tail, miRNA precursors can be subdivided into the following segments: the 5’ region, miR-5p, the loop region, miR-3p and the 3’ region (Figure 2A). sRNAs (18 to 34 nt) and tyRNAs (10 to 17 nt) mapped to these regions with different patterns of abundance (Figure 2B). In all of our samples, sRNAs originated almost exclusively from the miR-3p and miR-5p regions, which correspond to the mature miRNA/miRNA* duplex. tyRNAs, however, originated mostly from the loop region of the precursor, followed in abundance by the 5’ and 3’ regions (Figure 2B). These results suggest that MIRNA-derived tyRNAs are largely products of the remnant pieces following cleavage by DICER-LIKE 1 (DCL1). The terminal regions of miRNA precursors are known to undergo exonucleolytic decay via a process that includes RISC-INTERACTING CLEARING 3’-5’ EXORIBONUCLEASES (RICE1) to clear 5’ fragments (Zhang et al., 2017; Yu et al., 2017b) and EXORIBONUCLEASE 4 (XRN4) to clear 3’ fragments (Souret et al., 2004). However, the process by which the leftover loop region is degraded is less well examined, but apparently results in the production of abundant tyRNAs.

**Figure 2.**
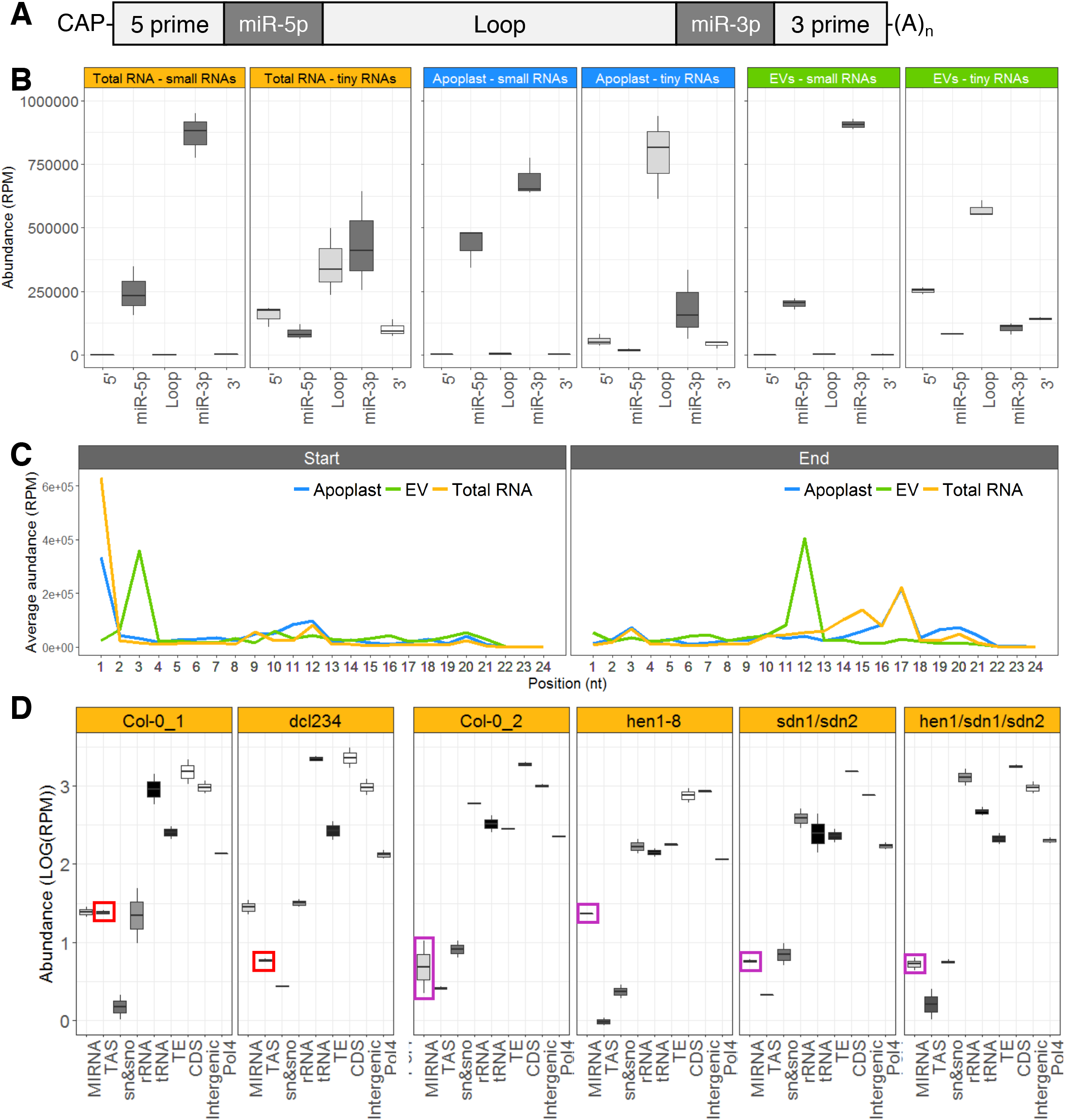
EV tyRNAs are Derived from Specific Regions of Precursor RNAs. **(A)** microRNA precursor structure, including from left to right, 5’ region, mature miRNA miR-5p, loop region, mature miRNA miR-3p, and 3’ region. **(B)** EV tyRNAs are enriched in sequences that map to the loop region of miRNA precursors. The abundance of small RNAs and tiny RNAs mapping to different regions of a miRNA precursor, as described in panel A. The y axis represents abundance, in reads per million (RPM). **(C)** EV tyRNAs mapping to mature miRNAs are cut out of the central region. Plots show the relative position of 5’ ends (left plot) or 3’ ends (right plot) of tyRNAs relative to nucleotide position within the parent miRNA. **(D)** Mutations that affect abundance of specific subclasses of sRNAs have a corresponding influence on tyRNA abundance. The abundance of total leaf tyRNAs mapping to different features of the *Arabidopsis* genome. These include the following from left to right: *miRNA* precursors, *TAS* precursors, snRNA and snoRNAs; ribosomal RNAs; tRNAs; transposable elements; coding sequences (including introns and UTRs); intergenic sequences; and known Pol IV precursors. The y axis represents abundance, in reads per million (RPM), in a logarithmic (LOG_10_) scale. Colored boxes highlight tyRNAs affected by mutations.

We also analyzed tyRNA abundances in a *dcl234* triple mutant, which is deficient for the production of 22-nt and 24nt siRNAs, including transacting siRNAs (tasiRNAs; (Henderson et al., 2006), but is unaffected in miRNA biogenesis. Consistent with expectations, MIRNA-derived tyRNA accumulation did not differ between the *dcl234* triple mutant and wild-type Col-0. However, we observed a clear reduction in TAS-derived tyRNAs (Figure 2C), indicating that these tyRNAs are likely derived from tasiRNA precursors.

Next, we examined data previously generated from *hen1, sdn1/sdn2* and *hen1/sdn1/sdn2* mutants (Yu et al., 2017a; samples included in the PRJNA251351 GEO dataset). HEN1 (HUA ENHANCER 1) is a methyl transferase that stabilizes specific sRNAs by adding a methyl group at the 3’ end, which protects them from degradation (Li et al., 2005). When HEN1 is absent, sRNAs undergo an uridylation process that allows exonucleases encoded by *SDN1* (*SMALL RNA DEGRADING NUCLEASE*) and *SDN2* to degrade them. We observed that MIRNA-derived tyRNAs increased in *hen1* mutants relative to wild-type Col-0, when miRNAs are destabilized, but remained at wild-type levels in sdn1/sdn2 and *hen1/sdn1/sdn2* mutants, which lack the exonucleases responsible for miRNA degradation (Figure 2C). These results support our hypothesis that MIRNA-derived tyRNAs are degradation products.

tyRNAs also mapped to the miRNA/miRNA* duplex, and we next checked the position of the tyRNAs within these ~21 or 22 nt molecules (Figure 2D). Since the miRNA and miRNA* designations were removed from miRBase, we treated both sides of the duplex equally. The majority of EV tyRNAs that mapped to miRNAs originated from the center of the miRNA sequence (i.e. nucleotides 3 to 12) (Figure 2D). In contrast, tyRNAs from total leaf libraries and apoplast mostly started at position 1 of the miRNA, followed by position 11, and they have an end position more widely distributed, from positions 13 to 21. These observations suggest that EV tyRNAs are not a random sampling of total cell tyRNAs, but are instead derived from a specific packaging process.

### 1-hit tyRNAs map to coding CDSs and intergenic regions

Due to the short length of tyRNAs, most tyRNAs map to multiple locations in the genome. To better determine their origin, we repeated the previously described analyses using only reads that map once to the genome (hereafter “1-hit” reads). These 1-hit tyRNAs have a different size distribution, ranging from 13 to 17 nt long, with the majority being 16 to 17 nt (Supplemental Figure 1A).

The majority of the 1-hit tyRNAs mapped to coding (CDS) and intergenic regions (Figure 3A). We examined in more detail the exact origin of the CDS-derived tyRNAs by dividing each gene into 5’UTR, exons, introns and 3’UTR. The 1-hit tyRNAs originated mostly from the sense strand of exons (Figure 3B), with no difference regarding the size of the tyRNAs (Supplemental Figure 1B). These results were independent of the sample type, suggesting that most of the CDS-derived tyRNAs originate from mature mRNAs.

**Figure 3.**
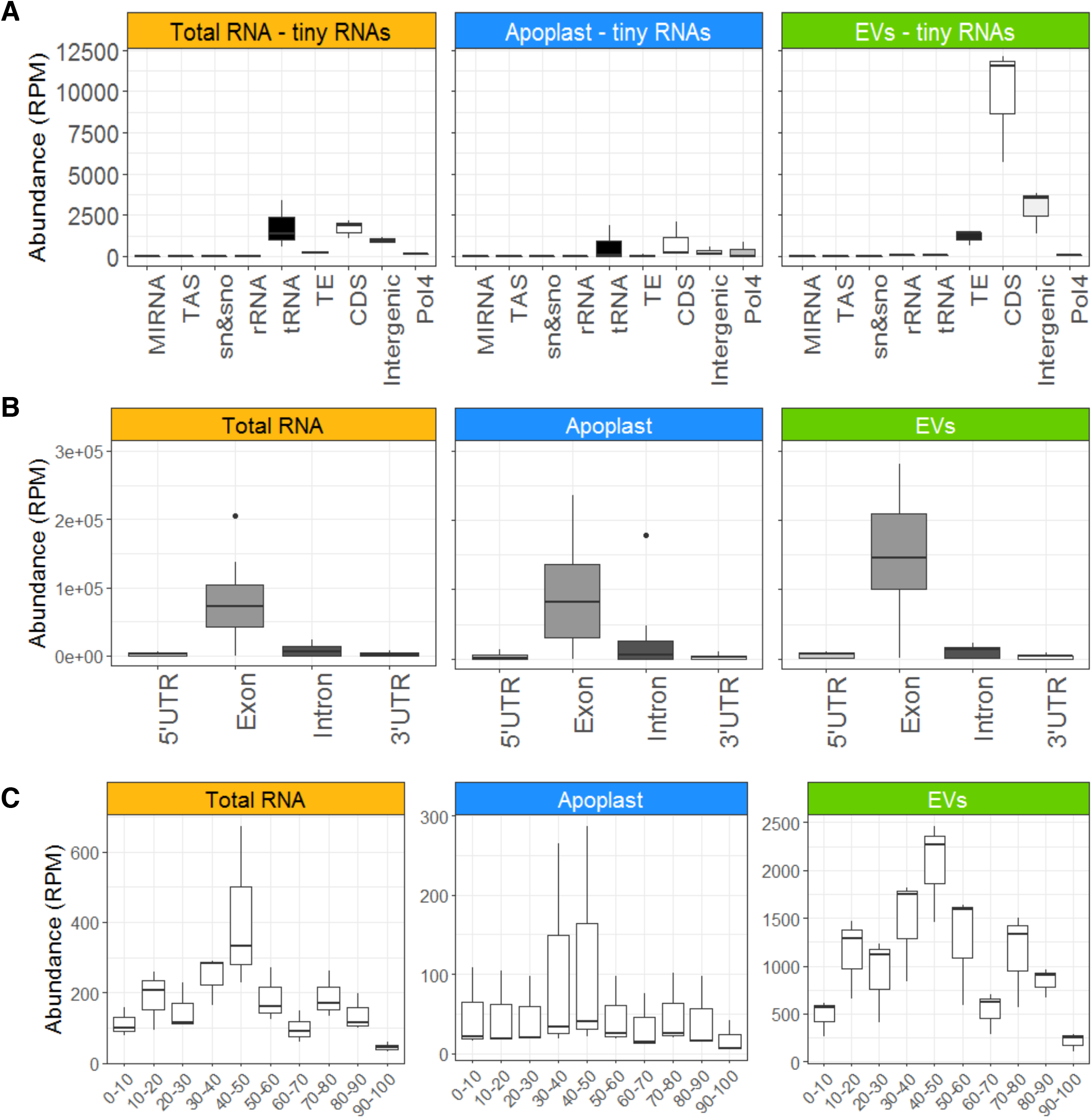
tyRNAs that Map to a Single Position in the Genome are Enriched in Exon-Derived Sense-Strand Sequences. **(A)**The abundance of 1-hit tyRNAs mapping to different features of the *Arabidopsis* genome. **(B)** The abundance of 1-hit tyRNAs mapping to 5’UTRs, exons, introns, 3’UTRs and intergenic regions. The y axis represents abundance, in reads per million (RPM) **(C)** The abundance of 1-hit tyRNAs mapping to different relative positions in mRNAs. The x axis represents the relative position in full-length mRNA, expressed in percentage of the total length, and the y axis represents abundance, in reads per million (RPM). Note that scales are different for each plot.

To further characterize the origin of the exon-derived tyRNAs, we analyzed their 5’ positions relative to the full-length mRNA, expressed as a percentage of the total length. In all samples, we observed a consistent peak originating at the center of the mRNA (Figure 3C). This persistent peak was more prominent in the EV samples than in whole leaf and apoplast samples. This pattern suggests that mRNAs may be first degraded by 5’ and 3’ exonucleases, with tyRNAs being derived from what is left over.

### EVs preferentially load specific miRNAs

We next investigated whether sRNAs accumulate differentially in EVs relative to the apoplast and total leaf samples, focusing on three major classes of sRNAs (heterochromatic siRNAs or hc-siRNAs, phasiRNAs, and miRNAs) (Figure 4). The most differentially represented sRNAs were miRNAs (Figure 4A). Group I consisted of miRNAs that were more abundant in EVs and apoplast samples compared to total RNA. While the miRNAs within this group seem to be secreted from the cell, there is no specific accumulation in EVs. Group II miRNAs accumulated to high levels in EVs and were present at slightly lower levels in the apoplast samples, suggesting some accumulation in EVs. The miRNAs in group III were depleted in the apoplast samples relative to total RNA and EVs, suggesting that group III RNAs are preferentially secreted via EVs. The miRNAs in group IV had an opposite accumulation pattern, low in EVs compared to apoplast and high in apoplast compared to total RNA samples. These results indicate that a distinct subset of miRNAs are preferentially loaded into the EVs.

**Figure 4.**
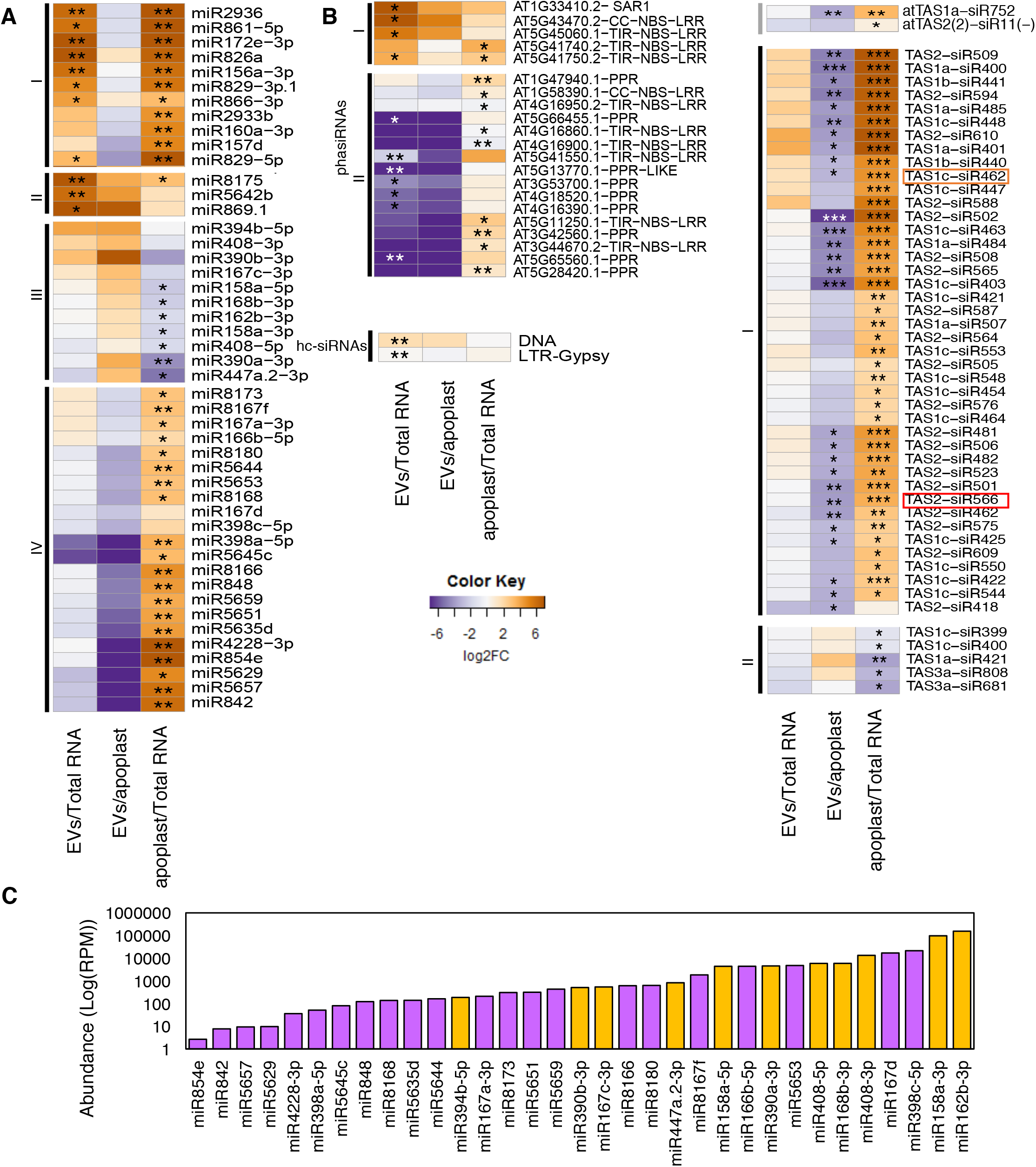
A Subset of Small RNAs Display Differential Abundance in EVs Versus Total leaf RNA or Apoplastic RNA. **(A)** Select miRNAs are enriched or depleted in EVs. Relative abundance of miRNAs in EVs and apoplast compared to abundance in total leaf RNA; the relative abundance is expressed as a heat map (see key on lower right), with the samples being compared indicated below each heat map (*q-value,≤0.05 and **q-value≤0.01). **(B)** Select siRNAs are enriched in EVs. Heat map indicates relative abundance of siRNAs (including phasiRNAs, tasiRNAs and hc-siRNAs) in the indicated samples. Boxed tasiRNAs indicate sRNAs shown to be taken up by *Botrytis cinerea* in Cai *et al*., 2018 (*q-value≤0.05, **q-value≤0.01 and ***q-value ≤0.001). **(C)** miRNA abundance does not correlate with EV enrichment. Absolute abundance in EVs of select miRNAs expressed in reads per million (RPM) in logarithmic scale, including miRNAs enriched in EVs (orange) and those depleted in EVs (purple) relative to apoplastic RNA.

The four patterns of preferential miRNA secretion described above did not correlate with total miRNA abundance in leaves. miRNAs with similar abundances in the total leaf sample were represented in both groups III and IV (i.e. those that accumulated to both low and high levels in the EV samples; Figure 4C). For example, miRNAs miR390b-3p, miR167c-3p, miR8166 and miR8180 had a similar abundance in total RNA samples, but the first two were abundant in EV samples while the latter two displayed a low abundance in EVs.

Combined, our findings hint at two separate pathways for the secretion of miRNAs. While miRNAs present in groups I and II may be secreted through either pathway, with a slight preference for EV-dependent secretion in group II, miRNAs in group III are strongly dependent on EVs for secretion. In contrast, miRNAs in group IV are secreted primarily through an EV-independent pathway. There may be specific sorting mechanisms for loading miRNAs into these separate pathways. Whether such specificity might be directed by the miRNA sequence or duplex structure, or something about its target or function remains unknown.

### siRNAs are poorly represented in EVs

We next assessed the abundance of full-length siRNAs in EVs, including phased siRNAs (phasiRNAs), tasiRNAs and heterochromatic siRNAs (hc-siRNAs). For phasiRNAs, we analyzed all members of families of genes encoding nucleotide-binding site leucine-rich repeat proteins (NLRs), pentatricopeptide repeat proteins (PPR) and SUPPRESSOR OF AUXIN RESISTANCE proteins (SAR), which are the primary sources of phasiRNAs in *Arabidopsis* (Supplemental Table 2). We were able to identify 21 phasiRNA producing loci that were differentially accumulated in one of our three comparisons (Fig. 4B). These loci can be divided into two groups; group one had high accumulation in EV and apoplast samples, compared to total RNA, and group II had a low accumulation in EV samples compared to apoplast and total RNA samples, and a high accumulation in apoplast compared to total RNAs. Similarly, we analyzed the accumulation of the 24 different *Arabidopsis* tasiRNAs that have been identified (Zhang et al., 2014) (Supplemental Table 2), and found only two that were differentially accumulated (Figure 4C – upper panel), with low accumulation levels in EVs. We also analyzed 21- and 22-nt RNAs derived from the eight *TAS* genes identified in the *Arabidopsis* genome for their specific differential accumulation in our samples (Figure 4C – lower panel). We observed that none of these TAS-derived sRNAs accumulated in EVs. Instead, all of the differentially accumulated sRNAs preferentially accumulated in the apoplast. This may indicate that phasiRNAs and tasiRNAs are preferentially directed to an apoplastic secretion pathway that is independent of EVs.

hc-siRNAs largely originate from transposable elements (TEs) and are thus repetitive in their nature, confounding analyses that require knowing their exact origin. Some hc-siRNAs may map to more than one TE with all the mapping locations corresponding to a single TE superfamily. Using hc-siRNAs that mapped within single superfamilies (Supplemental Table 2 - Raw reads all sRNAs), we measured hc-siRNA levels and observed no clear patterns of accumulation in EV samples. Therefore, unlike miRNAs that demonstrated specificity in their EV accumulation, there was no evidence of specificity in EV-localized sRNAs from loci generating either hc-siRNA or any other siRNA analyzed here.

## CONCLUSIONS

Here we have shown that EVs are highly enriched in small RNAs 10 to 17 nucleotides in length, which we have named tiny RNAs. tyRNAs appear to be degradation products derived from multiple sources, including mRNAs, primary miRNAs, siRNAs, tasiRNAs and hcRNAs. Whether this new class of RNAs have a cellular function, or represent a waste-or by-product of cellular metabolism, remains unknown.

One potential function of tyRNAs could be as small activating RNAs (saRNAs). RNA activation is a mechanism by which small RNAs, usually 18 to 21 nt long, can activate gene transcription, and was first identified in human cells more than a decade ago (Li et al., 2006). More recently, several studies have identified animal saRNAs that are able to positively regulate gene expression by targeting UTRs (Vasudevan and Steitz, 2007; Rocha et al., 2007; Ørom et al., 2008). Since then, more than 15 examples of saRNAs have been described in animal model systems, such as rats and primates, *in vivo* and *in vitro* (Huang et al., 2010). saRNAs bind to promoter regions in a wide range of positions relative to their transcription starting site (TSS) (Li et al., 2006; Janowski et al., 2007), as well as the 3’ UTR (Yue et al., 2010).

Here we also described that some miRNAs are specifically loaded into EVs, which reinforces the theory that sRNAs use EVs for long distance movement through the plant, and possibly as a cross-kingdom delivery system. Recently it has been described that plants secrete EVs to transfer sRNAs into fungal pathogens (Cai et al., 2018). Specifically, it has been shown that two plant-derived tasiRNAs from *TAS1c* and *TAS2* genes are transferred to *Botrytis cinerea* via detergent-sensitive particles to target genes involved in vesicle trafficking. We also found that some TAS1- and TAS2-derived siRNAs are secreted out of the cell, but mainly through an apoplastic secretion mechanism. The discrepancies between our findings and those of Cai et al. (2018b) may be due to differences in isolation methods, as Cai et al. used 100,000 x g as the relative centrifugal force rather than 40,000 x g to isolate EVs and they did not purify the vesicles away from any contaminating protein complexes. It may be that the tasiRNAs detected in Cai et al. were associated with other extracellular particles separate and distinct from EVs. This makes sense if, as our data suggests, plants employ multiple mechanisms for the secretion and long-distance transport of sRNAs.

## METHODS

### Plant materials and growth conditions

*Arabidopsis* seeds were germinated on 0.5X Murashige and Skoog (MS) medium containing 0.8% agar as previously described (Rutter and Innes 2017). The seeds were stored at 4°C for two days and then moved to short day conditions (9 hour days, 22°C, 150 μEm^−2^s^−1^). The seedlings were transferred after one week to Pro-Mix B Biofungicide potting mix supplemented with Osmocote slow-release fertilizer (14-14-14).

### EV isolation and purification

*Arabidopsis* EVs were isolated from apoplastic wash and purified on an iodixanol density gradient, as previously described (Rutter and Innes 2017). Briefly, 6 week-old *Arabidopsis* rosettes were vacuum infiltrated with Vesicle Isolation Buffer (VIB; 20 mM MES, 2 mM CaCl_2_, 0.01 M NaCl, pH 6.0). The rosettes were gently blotted to remove excess buffer and packaged inside needless, 30 ml syringes. The syringes were centrifuged for 20 min at 700 x g (2°C, JA-14 rotor, Avanti^™^ J-20 XP Centrifuge, Beckman Coulter, Indianapolis IN). The apoplastic wash collected from the rosettes was then filtered through a 0.22 μm membrane and centrifuged successively at 10,000 x g for 30 min to remove any large particles, 40,000 x g for 60 min to pellet the EVs and again at 40,000 x g to wash and re-pellet the EVs (2°C, SW41, Optima XPN-100 Ultracentrifuge, Indianapolis IN). After the first 40,000 x g step, 5 mLs of the supernatant were retained on ice for RNA isolation. The washed EV pellet was resuspended in 1 ml of cold, sterile VIB and loaded on top of a discontinuous iodixanol gradient (5%, 10%, 20% and 40%) (Optiprep^™^; Sigma-Aldrich, St. Louis MO). The gradient underwent centrifugation at 100,000 x g for 17 hours (2°C, SW41, Optima XPN-100 Ultracentrifuge, Indianapolis IN). After centrifugation, the first 5 ml of the gradient were discarded. The next three fractions of 0.8 ml were collected. The fractions were diluted in cold, sterile VIB and re-pelleted at 100,000 x g for 60 min (2°C, TLA100.3, Optima^™^ TLX Ultracentrifuge, Beckman Coulter, Indianapolis IN). The pellets were resuspended in cold, sterile VIB, combined and brought up to a total volume of 100 μl using VIB.

### RNA Sample Preparation

RNA was isolated from three different sources: leaf tissue, EV-depleted supernatant and purified EVs. Leaf RNA was isolated from 100 mg of leaf tissue harvested after collecting apoplastic wash, frozen and ground to a powder using liquid nitrogen and suspended in 1 ml of TRIzol^™^ Reagent (Thermo Fischer Scientific, Waltham, MA). Supernatant was collected after the first 40,000 x g step, as described above, and combined with 0.1 volumes of 3 M sodium acetate and one volume of cold isopropanol. The supernatant sample was mixed briefly by vortexing and kept at -20°C for one hour. After one hour, the sample was centrifuged for 30 min at 12,000 x g and 4°C. The resulting pellet of RNA was resuspended in 1 ml of TRIzol^™^ Reagent. EV RNA was isolated from EVs purified on a discontinuous iodixanol gradient as described above. 100 μl of EVs was added to 1 ml of TRIzol^™^ Reagent. Once all three samples were in TRIzol^™^ Reagent, RNA was extracted following the manufacturer’s instructions.

### Small RNA libraries

Small RNA libraries were constructed as described (Mathioni et al., 2017) using the NEBNext^®^ Small RNA Library Prep Set (NEB #E7330S). Due to the low abundance of RNA present in extracellular vesicles and apoplast samples, 500 ng of total RNA was used as the starting material. Libraries were sequenced on a NextSeq instrument with single-end 75-bp reads at the Indiana University Center for Genomics and Bioinformatics (Bloomington, IN).

### Data analysis

Small RNA sequencing libraries were trimmed for adaptors using cutadapt v1.16 (Martin, 2011) with a minimum insert size of 10 nt and a maximum of 34 nt. Sequence quality was assessed using FastQC (http://www.bioinformatics.babraham.ac.uk/projects/fastqc/). Clean reads were aligned to the *Arabidopsis* genome (TAIR version 10), and all subsequent analysis were done using Bowtie2 (Langmead and Salzberg, 2012). For miRNA and tasiRNA analyses, the latest versions of miRBase (v22 - (Kozomara and Griffiths-Jones, 2014) and tasiRNA database (Zhang et al., 2014) were used, respectively. Mapping positions for miRNAs were assessed using the Bowtie2 output file. Differential accumulation analyses were performed using DEseq2 with default parameters, using not normalized reads as input (Love et al., 2014) and graphical representations using ggplot2 (Wickham, 2009) in the R statistical environment. For box and whisker plots produced by ggplot2, the upper whisker represents the maximum value, or the top of the box plus 1.5 IQR, whichever is less (IQR = the length of the box), while the lower whisker represents the minimum value, or the bottom of the box minus 1.5 IQR, whichever is greater.

### Micrococcal nuclease protection assays

EVs were isolated from 10 mL of apoplastic wash fluid collected from 36 *Arabidopsis* rosettes, as described above. The EV pellet obtained after the second centrifugation step at 40,000 x g was re-suspended in 250 μL of sterile VIB (pH 7.5) supplemented with 1X micrococcal nuclease reaction buffer (New England BioLabs; 50 mM Tris-HCl, 5 mM CaCl_2_, pH 7.9). An equal amount of re-suspended EVs (50 μL) was used for each treatment with or without membrane permeabilization prior to microccoal nuclease addition. To permeabilize EVs, 0.075% (v/v) Triton X-100 was added to EVs and vortexed for 1 minute. For nuclease treatments, 0.5 μL (1000 units) of micrococcal nuclease (New England BioLabs catalog number M0247S) was added and incubated at 37°C for 5 minutes. Digestion was terminated by addition of EGTA (final concentration: 20 mM) and RNA was immediately isolated using TRIzol reagent (Invitrogen) as described above. Both microRNAs and tyRNAs levels were quantified using stem-loop qRT-PCR as described by Varkonyi-Gasic et al. (2007), starting with same volume of sample from each treatment. Primers used for first strand cDNA synthesis and qRT-PCR are provided in Supplemental Table 3.

### Accession Numbers

All RNA sequencing data has been deposited into the Gene Expression Omnibus (GEO) under the accession code GSE114696.

## Supplemental Data

**Supplemental Figure 1**: Characterization of 1-hit tyRNAs.

**Supplemental Table 1**: small RNA sequencing summary statistics.

**Supplemental Table 2**: Raw read count by sRNA class.

**Supplemental Table 3**: Primers used for qRT-PCR analyses.

## ACKNOWLEDGMENTS

We thank Craig Pikaard for providing seed of *dcl2/3/4 Arabidopsis* triple mutant, and the Indiana University Physical Biochemistry Instrumentation Facility for access to ultracentrifuges and nanoparticle tracking equipment. We also thank the IU Center for Genomics and Bioinformatics for assistance with the generation and analysis of sRNA-seq data. This work was supported by three grants from the United States National Science Foundation, IOS-1645745 and IOS-1842685 to RWI and IOS-1842698 to BCM.

## AUTHOR CONTRIBUTIONS

P.B., B.D.R., R.W.I. and B.C.M. designed the research. P.B., B.D.R. and H.Z. performed the research. P.B. and R.P. analyzed the data. P.B., B.D.R. and R.W.I. wrote the article. All authors read and commented on the manuscript.

